# Attentional bias toward fearful faces is associated with maternal postnatal distress and alterations in white matter microstructure in 5-year-old females

**DOI:** 10.1101/2023.11.01.565103

**Authors:** Venla Kumpulainen, Eeva-Leena Kataja, Elmo P. Pulli, Anni M. Copeland, Eero A. Silver, Tuomo Haikio, Ekaterina Saukko, Riikka Korja, Linnea Karlsson, Hasse Karlsson, Jetro J. Tuulari

## Abstract

The development of face processing system is paramount for social interaction and communication skills. Eye-tracking provides a viable means to investigate face processing even in very young children and can be used to derive indices of affective processing biases that are adaptive in supporting preparedness for emotional encoding. The neural correlates of early-emerging attentional biases have not been explored.

In the current study, we gathered diffusion tensor imaging (DTI) and eye-tracking data (attention disengagement from neutral, happy, fearful faces, and control pictures towards distractor stimuli) to investigate the association of white matter (WM) microstructural alterations and attentional biases in 117 (55 female) typically developing 5-year-old children. We estimated fractional anisotropy (FA) and used tract-based spatial statistics (TBSS) before conducting voxel-wise analyses testing associations between attentional biases and brain diffusivity.

We found that reduced WM integrity (indexed as decrease in FA) in widespread WM regions, including the splenium of corpus callosum, left anterior limb of internal capsule (ALIC), left posterior limb of internal capsule (PLIC), left posterior thalamic radiation and optic tract, predicted higher attentional bias toward fearful expressions in females (adjusted for potential confounders). Further, maternal postnatal anxiety and depressive symptoms were detected to associate positively with attentional bias toward fearful expression, but only in females.

Based on these findings and prior results on association between maternal distress and reduced WM integrity in female offspring, it is possible that alterations in WM microstructure may transmit the long-term effects of maternal mental distress during early life to increased vigilance towards negative emotional expressions. These hypotheses and sexual dimorphism in the observed associations require replication and are potential focus areas for future studies.

## 1 Introduction

The development of face processing system is paramount for social interaction and communication skills. Face present prompt cues of the identity, intentions, and the emotional state of the person. The recognition of faces develops early, as infants have been shown to prefer viewing faces versus other objects (Johnson, 2007), and the ability to identify a person by facial features is developed already by the age of 3 to 5 months (Bhatt et al., 2005; Hayden et al., 2007). The emotional facial expressions are essential non-verbal means of social interaction and for sharing the feelings or the emotional states between individuals. The processing of facial expressions includes the identification of emotional cues, reacting with an adequate affective state and regulating the emotional behaviour with situational demands (Phillips et al., 2003). Early social interaction with parents and parental behaviour seem to model the neural responses to emotional faces (Taylor-Colls & Pasco Fearon, 2015). Maternal postnatal depressive symptoms have been shown to predict increased attention toward fearful emotional expression in infants (Forssman et al., 2014; E.-L. Kataja et al., 2020), and these cognitive biases toward fear and threat may be transmitted from mother to offspring (Fliek et al., 2017; Remmerswaal et al., 2016). Further, vigilance toward threat signals in children has been associated with increased risk for anxiety (Abend et al., 2018) and to later avoidance of threat (Remmerswaal et al., 2016).

The core face processing areas of the brain include the occipital face area (OFA), fusiform face area (FFA) and posterior superior temporal sulcus (pSTS) (Haist & Anzures, 2017). Activation of OFA and FFA have been associated with general identification of faces, while pSTS participates in processing of more dynamic and changing characteristics like facial expressions (Isik et al., 2017). The extended face system, which includes shared regions recruited in emotional processing generally, participates in interpretation of social and emotional contexts related to the face recognition and is recruited in a task-specific manner especially in adults (Haist et al., 2013). Literature of neural correlates in emotional processing has not revealed one specific brain region recruited in variable emotional tasks. However, multiple areas have been shown to activate consistently with emotional stimuli. Among these structures, limbic regions (Gobbini & Haxby, 2007; Ishai et al., 2004), the amygdala and its connections, prefrontal cortex (PFC), orbitofrontal cortex (OFC) and anterior cingulate cortex (ACC) are pivotal in both the emotional perception and regulation, and they develop in divergent schedule with the amygdala maturing earlier in the childhood in contrast to PFC and ACC maturing into the late adolescence (Pechtel & Pizzagalli, 2012; Qin et al., 2012). Fronto-limbic connectivity changes during childhood and adolescence from positive to negative, and presumably increases the prefrontal regulation of the amygdala reactivity (Gabard-Durnam et al., 2014).

Main connections of the face processing areas travel through inferior longitudinal fasciculus (ILF) and inferior fronto-occipital fasciculus (IFOF) connecting the frontal/temporal and the occipital regions. While ILF participates in face recognition (Catani et al., 2003; C. J. Fox et al., 2008), visual perception (Ffytche & Catani, 2005) and visual memory (Ross, 2008), IFOF has been shown to be particularly important for visual processing (C. J. Fox et al., 2008), attention (Doricchi et al., 2008) and recognition of emotional facial expressions (Philippi et al., 2009). Additionally, direct connections between OFA and FFA (but not with STS), and between visual areas and the amygdala have been detected with tractography with predominance in the right hemisphere (Gschwind et al., 2012). This is in accordance with the previous notions suggesting right-sided dominance in face processing (C. J. Fox et al., 2008). The pSTS, on the other hand, has been shown to have connections to more anterior temporal, superior parietal, and frontal regions in response to face perception (Gschwind et al., 2012).

The behavioural studies have provided evidence of infants’ ability to categorize facial emotional expressions between the ages of 4 and 8 months (Barrera & Maurer, 1981; Caron et al., 1982; Nelson, 1987). While general preference for faces and attentional bias toward basic emotional expressions is detected in infants, the emotional understanding is developing only later in childhood (Denham et al., 2003; C. M. Herba et al., 2006). Infants show bias toward looking at happy faces as early as at age of 4 to 6 months (Farroni et al., 2007; LaBarbera et al., 1976). However, the attentional bias shifts from positive to negative emotional expressions, especially to fearful faces, already during the first year of life (Vaish et al., 2008). This shift has been suggested to reflect the transition of completely dependent infant, requiring all resources from a caregiver to a more mobile and autonomic individual with more needs to be able to avoid potential harms (Elam et al., 2010). The attentional bias to both happy and fearful faces is detected again among 5-year-olds (Elam et al., 2010), suggesting age-related changes in face and fear processing patterns across early childhood. In a previous study of the current cohort (E.-L. Kataja et al., 2022), attentional disengagement from faces to a distractor was observed to increase from infancy (8 months) to toddlerhood (30 months), but similar trend was not observed when the results were compared to children aged 60 months, thus reflecting earlier findings (Libertus et al., 2017). This implies a U-shaped trajectory of attentional disengagement pattern during development. Further, the recognition of emotional expressions follows different developmental trajectories in males and females (Mancini et al., 2013). For example, females showed more accurate recognition of sad facial expression at the age of 8 years, yet males closed the gap by the age of 11 years (Mancini et al., 2013). Currently, little is known about the anatomical pathways underlying face and fear biases during early childhood.

Affective processing biases are adaptive in supporting preparedness for fear responses. However, aberrant stress reactivity, including excessive attentional bias to threat, is associated with increased internalizing symptoms (Carlson et al., 2013, 2014) and anxiety, also in pediatric populations (Abend et al., 2018) (Table 1 summarizes previous studies of attentional bias to threat or fear stimuli in pediatric populations). Among normative adult population, Carlson et al. (2013) found that greater attention bias to threat predicted greater levels of uncinate fasciculus integrity, greater positive amygdala and anterior cingulate functional connectivity, and greater amygdala coupling with a broader social perception network including the STS, temporoparietal junction (TPJ), and somatosensory cortex. In another study, Carlson et al. (2014) showed a positive association between amygdalo-prefrontal white matter (WM) tract (UF) integrity and attention bias to threat. The study by Koller et al., (2019) showed higher FA of the superior colliculus (SC) and amygdala tract in the right hemisphere to predict stronger threat bias among healthy adults. However, the early neural correlates behind these features and observations have remained mostly elusive while this knowledge might increase understanding of early emerging individual differences in fear and threat sensitivity, for instance.

**Table 1.**
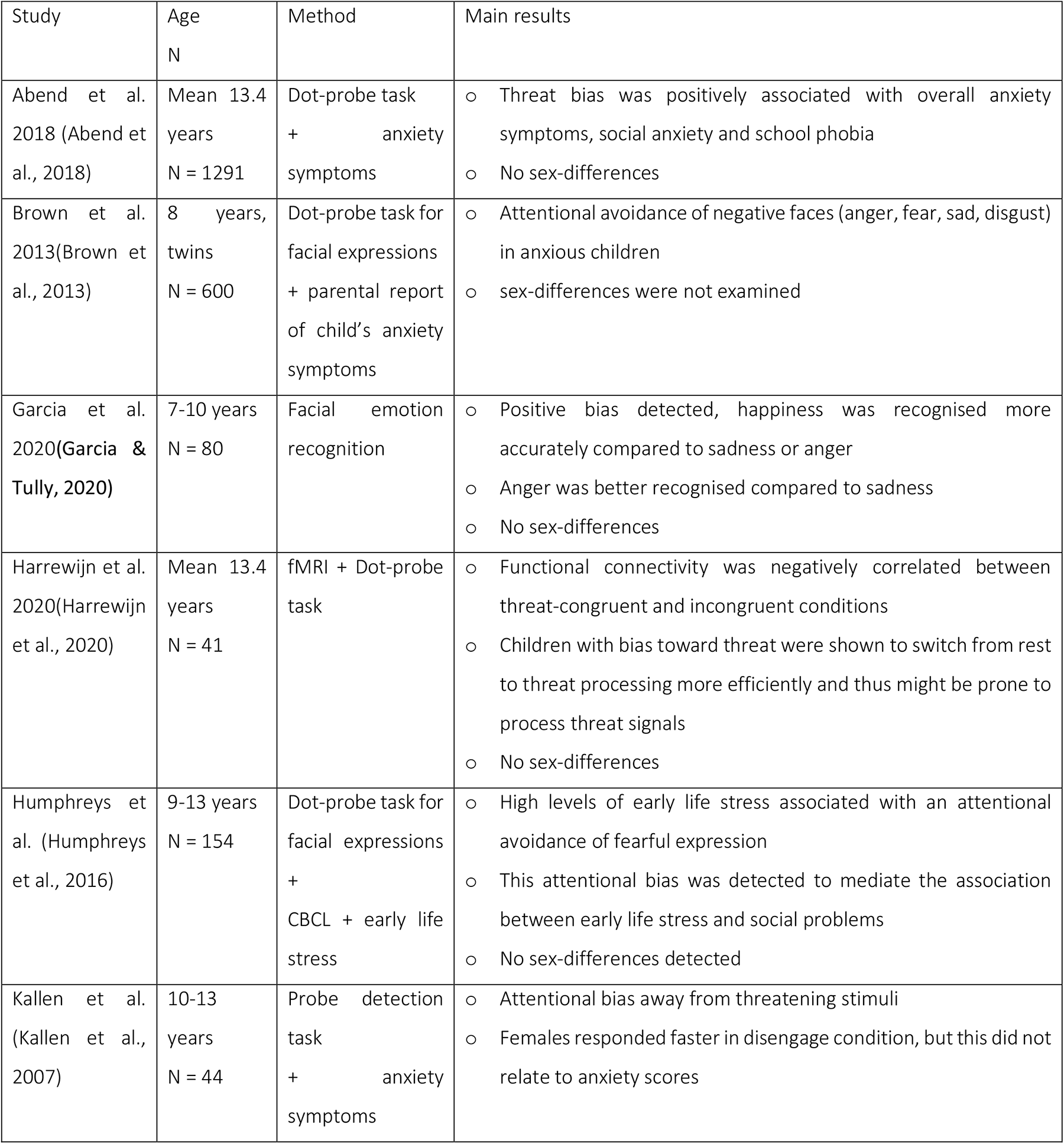
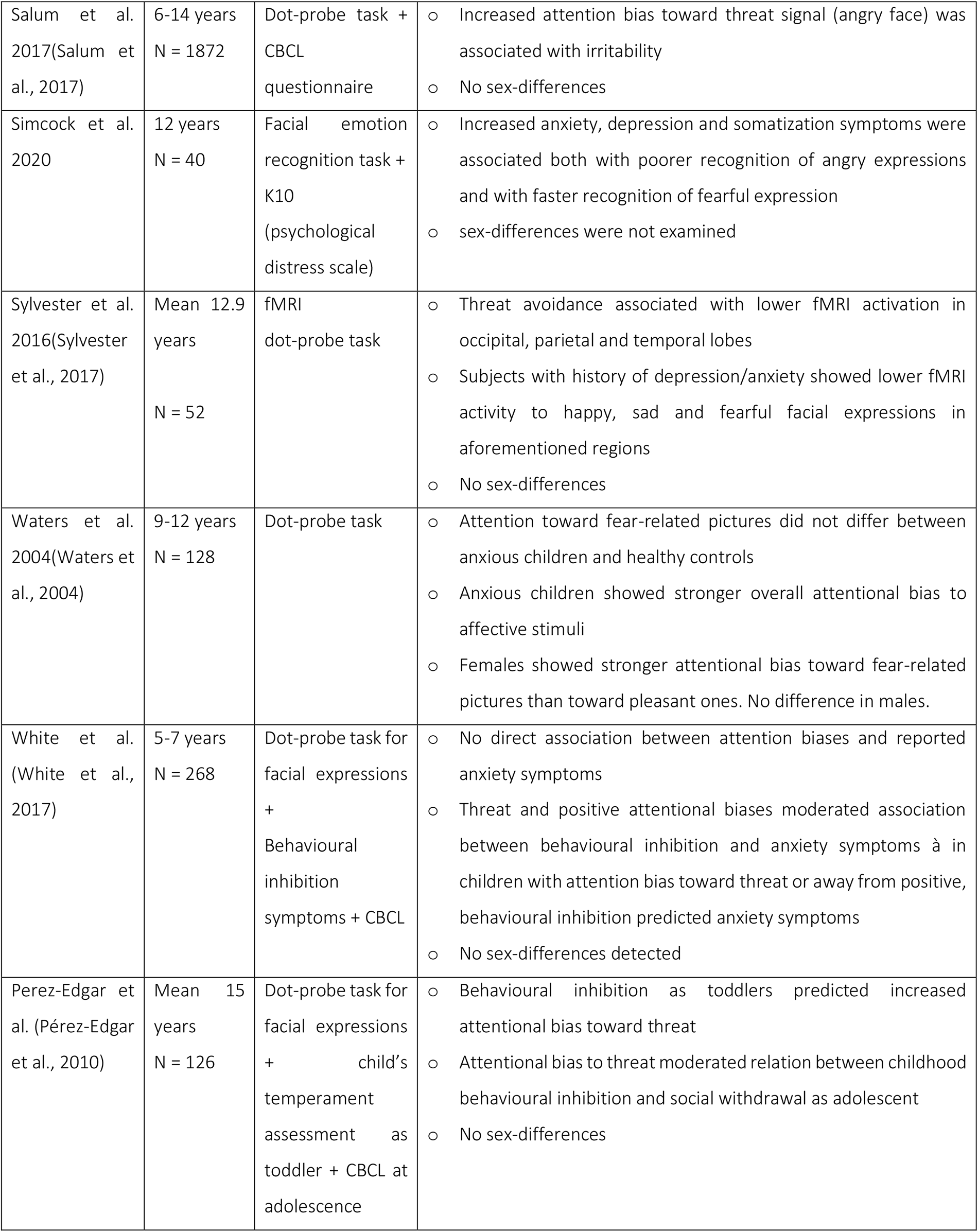
Literature review considering previous studies of attentional bias to threat or fear stimuli in pediatric populations.

| Study | Age<br>N | Method | Main results |
| --- | --- | --- | --- |
| Abend et al.<br>2018 (Abend et al., 2018) | Mean 13.4<br>years<br>N = 1291 | Dot-probe task<br>+ anxiety<br>symptoms | ○ Threat bias was positively associated with overall anxiety symptoms, social anxiety and school phobia<br>○ No sex-differences |
| Brown et al.<br>2013(Brown et al., 2013) | 8 years,<br>twins<br>N = 600 | Dot-probe task for<br>facial expressions<br>+ parental report<br>of child's anxiety<br>symptoms | ○ Attentional avoidance of negative faces (anger, fear, sad, disgust) in anxious children<br>○ sex-differences were not examined |
| Garcia et al.<br>2020(Garcia & Tully, 2020) | 7-10 years<br>N = 80 | Facial emotion<br>recognition | ○ Positive bias detected, happiness was recognised more accurately compared to sadness or anger<br>○ Anger was better recognised compared to sadness<br>○ No sex-differences |
| Harrewijn et al.<br>2020(Harrewijn et al., 2020) | Mean 13.4<br>years<br>N = 41 | fMRI + Dot-probe<br>task | ○ Functional connectivity was negatively correlated between threat-congruent and incongruent conditions<br>○ Children with bias toward threat were shown to switch from rest to threat processing more efficiently and thus might be prone to process threat signals<br>○ No sex-differences |
| Humphreys et al. (Humphreys et al., 2016) | 9-13 years<br>N = 154 | Dot-probe task for<br>facial expressions<br>+<br>CBCL + early life<br>stress | ○ High levels of early life stress associated with an attentional avoidance of fearful expression<br>○ This attentional bias was detected to mediate the association between early life stress and social problems<br>○ No sex-differences detected |
| Kallen et al.<br>(Kallen et al., 2007) | 10-13<br>years<br>N = 44 | Probe detection<br>task<br>+ anxiety<br>symptoms | ○ Attentional bias away from threatening stimuli<br>○ Females responded faster in disengage condition, but this did not relate to anxiety scores |
| Salum et al. 2017(Salum et al., 2017) | 6-14 years<br>N = 1872 | Dot-probe task + CBCL questionnaire | ○ Increased attention bias toward threat signal (angry face) was associated with irritability<br>○ No sex-differences |
| Simcock et al. 2020 | 12 years<br>N = 40 | Facial emotion recognition task + K10 (psychological distress scale) | ○ Increased anxiety, depression and somatization symptoms were associated both with poorer recognition of angry expressions and with faster recognition of fearful expression<br>○ sex-differences were not examined |
| Sylvester et al. 2016(Sylvester et al., 2017) | Mean 12.9 years<br>N = 52 | fMRI dot-probe task | ○ Threat avoidance associated with lower fMRI activation in occipital, parietal and temporal lobes<br>○ Subjects with history of depression/anxiety showed lower fMRI activity to happy, sad and fearful facial expressions in aforementioned regions<br>○ No sex-differences |
| Waters et al. 2004(Waters et al., 2004) | 9-12 years<br>N = 128 | Dot-probe task | ○ Attention toward fear-related pictures did not differ between anxious children and healthy controls<br>○ Anxious children showed stronger overall attentional bias to affective stimuli<br>○ Females showed stronger attentional bias toward fear-related pictures than toward pleasant ones. No difference in males. |
| White et al. (White et al., 2017) | 5-7 years<br>N = 268 | Dot-probe task for facial expressions + Behavioural inhibition symptoms + CBCL | ○ No direct association between attention biases and reported anxiety symptoms<br>○ Threat and positive attentional biases moderated association between behavioural inhibition and anxiety symptoms → in children with attention bias toward threat or away from positive, behavioural inhibition predicted anxiety symptoms<br>○ No sex-differences detected |
| Perez-Edgar et al. (Pérez-Edgar et al., 2010) | Mean 15 years<br>N = 126 | Dot-probe task for facial expressions + child's temperament assessment as toddler + CBCL at adolescence | ○ Behavioural inhibition as toddlers predicted increased attentional bias toward threat<br>○ Attentional bias to threat moderated relation between childhood behavioural inhibition and social withdrawal as adolescent<br>○ No sex-differences |

Social emotional processing of young subjects can be studied, for example, by measuring visual attention. Eye-tracking provides a viable means to investigate face processing and attentional biases even in very young children. Previous studies have shown vigilance toward or avoidance of emotional expressions to relate to, for example, anxiety (Abend et al., 2018; Brown et al., 2013; Waters et al., 2004), other internalizing symptoms (Simcock et al., 2020; Sylvester et al., 2017), and behavioural inhibition (Pérez-Edgar et al., 2010; White et al., 2017) in children and adolescents. One previous study of 8-17-year-old children has examined the differences in WM microstructural correlates of attention to emotional facial expressions between brain tumour patients and healthy controls (Moxon-Emre et al., 2020). Increased integrity in corpus callosum was associated with worse emotion regulation in brain tumour patients, while in healthy controls they observed no correlations between eye-tracking scores and WM microstructure. To our knowledge, no study has reported associations between WM microstructural properties and attention biases in healthy young children.

In the current study, we gathered diffusion tensor imaging (DTI) and eye-tracking data to investigate the association of microstructural alterations of the white matter and attentional biases in healthy 5-year-old children. This was an exploratory study as this is to our knowledge the first DTI study to examine brain WM features predicting emotional attention in healthy preschool-aged children, and hence, we did not place a hypothesis on the magnitude or direction of the associations.

### 2 Methods

This study was reviewed and approved by the Ethics Committee of the Hospital District of Southwest Finland ((07.08.2018) §330, ETMK: 31/180/2011), and performed in accordance with the Declaration of Helsinki. Written informed consent was signed by both parents for the magnetic resonance imaging (MRI) measurement and by one parent for the eye-tracking experiments. The child assent was assured prior to neuroimaging visits.

### 2.1 Participants

The study population was part of the FinnBrain Birth Cohort Study (www.finnbrain.fi) (Karlsson et al., 2017) that was established to prospectively study the effects of exposure to early-life stress to the development of child brain, behaviour, and health. The subpopulation was invited to brain MRI based on attendance on prior neuropsychological examination visits, including eye-tracking experiments. All participants were healthy, normally developing, Finnish children aged 5.1 to 5.8 years (mean 5.4, SD 0.11; 111 males, 92 females; 182 right-handed, 14 left-handed, 7 with no preference) at the time of MRI.

The following exclusion criteria were used: 1) birth before gestational week (gwk) 35, 2) major developmental disorder or trait, 3) other long-term diagnosis requiring constant contact to hospital, 4) sensory abnormalities (e.g., blindness or deafness), 5) use of daily regular medication (asthma inhalers and one exception with desmopressin medication were allowed), 6) head trauma that had required inpatient care (reported by parents). Additional, metallic ear tubes and other common MRI contraindications inhibited the imaging.

Demographical data including sex, age at MRI, gestational age at birth, weight and height, exposure to smoking, selective serotonin reuptake inhibitors (SSRI) or synthetic glucocorticoids during pregnancy, maternal pre-pregnancy body mass index (BMI) and maternal socioeconomic status (classified by educational level, low-middle-high) were gathered. Maternal and obstetric variables were obtained from the Finnish Medical Birth Register of the National Institute for Health and Welfare. Additionally, maternal depressive and anxiety symptoms were surveyed during pregnancy (1^st^, 2^nd^ and 3^rd^ trimester) and 3, 6 and 12 months postpartum with Edinburgh Postnatal Depressive Scale (EPDS) (Cox et al., 1987) and Symptom Checklist-90 (SCL-90; not collected at 12 months postpartum) (Derogatis et al., 1973; Holi et al., 1998), respectively.

### 2.2 MRI visits

The MRI scans of 203 5-year-olds were performed between 29th October 2017 and 1st March 2021 in Turku University Hospital, Finland. The MRI was preceded by careful preparation including a home visit by a study nurse, home practicing period and training in hospital before MRI scanning. The children were imaged awake, and a staff member and parent(s) stayed in the scanner room through the imaging. Both ear plugs and noise cancelling headphones were used for hearing protection, and the child was also able to communicate through headphones and microphone. The imaging was ceased whenever the child showed signs of discomfort/wanted to discontinue, and taking breaks was also possible. The maximum length of the MRI scan was 60 minutes. Please see more detailed description of the MRI visit in our previous studies (Copeland et al., 2021; Kumpulainen et al., 2022; Pulli et al., 2022).

### 2.3 MRI sequences

The MRI scans were acquired with Siemens Magnetom Skyra fit 3T scanner (20-element head/neck matrix coil) and the Generalized Autocalibrating Partially Parallel Acquisition (GRAPPA) technique was used. DTI protocol was applied with a standard twice-refocused Spin Echo-Echo Planar Imaging (SE-EPI) and the following parameters: TR (time of repetition)/TE (time of echo) = 9300/87.0 ms, FOV (field of view) = 208 mm, isotropic voxels with 2.0x2.0x2.0 mm resolution, b value 1000 s/mm^2^, 96 noncollinear diffusion gradient directions divided into three scanning sets (31, 32 and 33 directions) with 9 b0 images (3 b0 = 0 s/mm^2^ volumes scattered within each set).

### 2.4 Pre-processing of diffusion data

Diffusion data was pre-processed with FSL 6.0 (FMRIB software library, University of Oxford, UK) including co-registration and averaging of b0 images (FLIRT, FMRIB’s Linear Image Registration Tool), creation of brain mask (BET, Brain Extraction Tool, 1.0.0; settings -R -f 0.3)(Smith, 2002), motion and eddy current correction and concurrent rotation of the b matrix. Prior to co-registration, DTIprep (https://nitrc.org/projects/dtiprep/ (Oguz et al., 2014)) and manual quality control were used to exclude volumes with motion artefacts and 30 directions per each subject was selected by maximizing the angular resolution (see prior studies considering validation of our pre-processing protocol (Kumpulainen et al., 2022; Merisaari et al., 2019)). The scalar maps were computed with FSL’s dtifit. Tract-based spatial statistics (TBSS) pipeline of FSL (Smith et al., 2006) was used to estimate the WM tract skeleton (tbss_2_reg flag -n, study-specific template, tbss_3_postreg flag -S, FA threshold 0.20). Tract-wise FA values were extracted with TBSS skeleton co-registration to JHU-ICBM-DTI atlas (Mori et al., 2008).

### 2.5 Eye-tracking measurements

The overlap paradigm (Peltola et al., 2009) was used to assess the child’s attention disengagement from faces and nonfaces to distractors (see Supplement Figure 1). In this paradigm, first, a picture of a neutral, happy, or fearful face or a nonface control stimulus was shown in the center of the screen for 1,000 ms. Then, a salient lateral distractor (i.e., a geometric shape), which started to flash after the child directed attention toward it, was added on either the left or right side of the screen (at visual angle of 13.6°) for 3,000 ms. One trial lasted for 4,000 ms. Altogether 24 pictures of one female model posing neutral, happy, and fearful faces and a nonface control stimulus were used.

The sizes of the central face/nonface stimuli and the lateral distractor stimuli were 15.4° 3 10.8° and 15.4° 3 4.3°, respectively. A brief animation was shown before each trial to capture the attention of the child to the center of the screen. Once the child was gazing in the middle of the screen, the trial was started manually by the researcher who monitored the child’s gaze from the host computer. The order of the central stimuli was semi-randomized, with a constraint that the same stimulus was not presented more than three times in a row. The lateral stimulus (i.e., black and white checkerboard pattern or vertically arranged empty and filled circles) was selected and presented randomly for each trial.

The eye-tracking data was pre-processed similarly to previous studies (E.-L. Kataja et al., 2022; Leppänen et al., 2015). The following criteria were used to ensure the quality of the trials retained in the analysis. These parameters were set a priori based on prior studies using the same methodology and analytic approach (Kataja et al., 2020; Leppänen et al., 2015). First, trials had to have sufficient looking at the central stimulus (> 70%) during a time interval that started at the onset of the trial, that is, appearance of the face or nonface on the screen, and extended to the end of the analysis period, that is, gaze disengagement from the central to lateral stimulus or if a gaze disengagement was not observed, 1,000 ms after the appearance of the lateral distractor. Second, trials had to have enough valid samples in the gaze data with no gaps greater than 200 ms. This meant that the gaps in the data were extrapolated by the analysis script by carrying the last recorded sample forward, but if the gap was greater than 200 ms, the trial was flagged as invalid and excluded from subsequent analyses. Third, if disengagement occurred during the trial, the exact timing of the eye movement from the central to the lateral stimulus was required to be known for the trial to be included in the analysis, that is, if the eye movement occurred during a period of missing or extrapolated gaze data, the trial was rejected.

In prior studies using the current methodology to assess child’s attentional disengagement, trial-level eye-tracking data have been analysed by coding the data into a binary disengagement value (0/1) based on whether or not the gaze shifted from the central to the lateral stimulus (i.e., disengagement probability, DP) or by extracting the latency of the gaze shift. The trial-level data have been aggregated to estimate mean DP or mean disengagement time (DT). The latter has also been variably calculated excluding or including censored values (i.e., trials on which no disengagement was observed by the end of the trial period (Pyykkö et al., 2019)). DPs and DTs are typically highly positively correlated, and the correlations were high in the current study as well (r_s_ = .89). In the current study, we used mean DPs as our primary measures and report these in the manuscript. Further, we calculated neutral/happy/fear bias indices (labelled as NE-bias, HA-bias, FE-bias, respectively) contrasting the DPs in mentioned face condition to DPs in nonface control condition (see Kataja et al., 2022). See Supplementary Table S1 for eye-tracking measures.

### 2.6 Statistical analysis

Statistical analyses with tabular data were performed with SPSS version 27.0 (IBM Corp. 2020, Armonk, NY, USA). First, we conducted group comparisons between females and males to determine whether they differed with respect to demographical variables, maternal distress scores or eye-tracking measurement results. Two-sample independent t-test was used for continuous variables and Chi-square test for categorical variables. Statistically significant differences in DPs and attentional biases between different facial expressions were examined with analysis-of-variances (ANOVA), and post-hoc Tukey’s HSD test (α level 0.05). Comparison between females’ and males’ attentional biases to different facial expressions was performed with paired sample t-test. Correlations between demographical variables, maternal distress scores and eye-tracking measures were examined with Spearman rank order correlation.

Voxel-wise analysis of scalar maps testing association between FA values and eye-tracking variables was performed with General Linear Model (GLM, FSL’s randomise tool). The analyses were performed first with all subjects and then for females and males separately, with 5000 permutations and multiple comparison correction with threshold-free cluster enhancement (TFCE). The following potentially confounding factors were included in the analyses: sex, child’s age at scan, maternal pre-pregnancy BMI, maternal age, exposure to smoking, SSRI or synthetic glucocorticoids (SGC) during pregnancy, maternal 2^nd^ trimester and 3 months postpartum EPDS and SCL-90 scores since early postnatal maternal distress was associated with WM metrics of interest (Kumpulainen, Copeland, et al., 2023).

## 3 Results

### 3.1 Demographics

A final sample of 117 subjects (55 females) with both eye-tracking and DTI data with sufficient quality after pre-processing steps was included in the study. Statistics for demographical data and eye-tracking variables are presented in Table 2. No statistically significant differences between females and males were observed in any of the background variables or eye-tracking measurements.

**Table 2.**
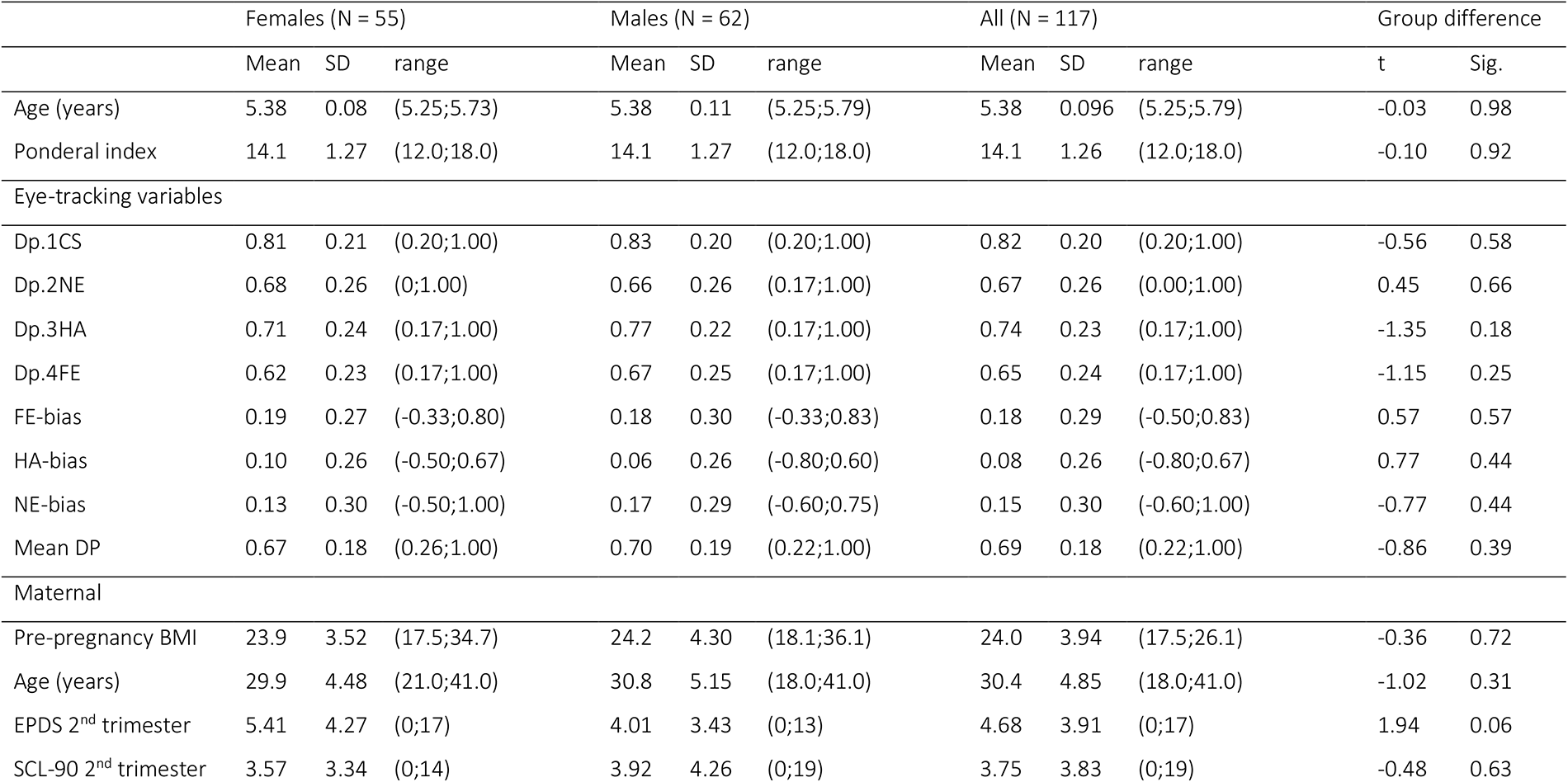
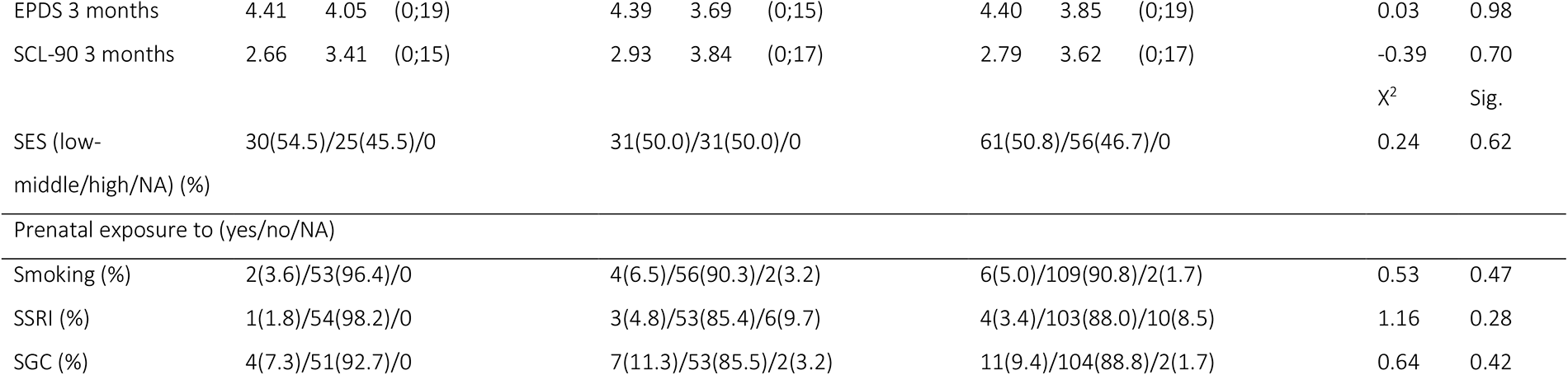
Demographical information, maternal psychological distress scores and eye-tracking measurements in females, males, and all subjects. Group differences between females and males examined with independent t test and chi-square test. SD = standard deviation, dp = disengagement probability, CS = control, NE = neutral, HA = happy, FE = fearful, EPDS = Edinburgh Posnatal depressive symptoms, SCL = Symptom Checklist, NA = not available, SES = socioeconomic status, SSRI = selective serotonin re-uptake inhibitor, SGC = synthetic glucocorticoid

### 3.2 Eye-tracking measurements

Disengagement probability was highest for control stimulus (0.82, SD 0.20; Table 2). Disengagement from neutral (0.67, SD 0.26, p < 0.001; Table 3), fearful (0.65, SD 0.24, p < 0.001) and happy (0.74, SD 0.23, p = 0.042) facial expressions were statistically significantly lower than in the control condition, except in males’ difference between disengagements from happy expression and control picture did not reach statistical significance (p = 0.059). Disengagement probability was higher for happy compared to fearful faces (mean difference 0.092, p = 0.014). Males were observed to disengage more likely from happy than neutral faces, too (mean difference 0.11, p = 0.002). No significant differences between females’ and males’ disengagement probability scores were observed with independent samples t test.

**Table 3.**
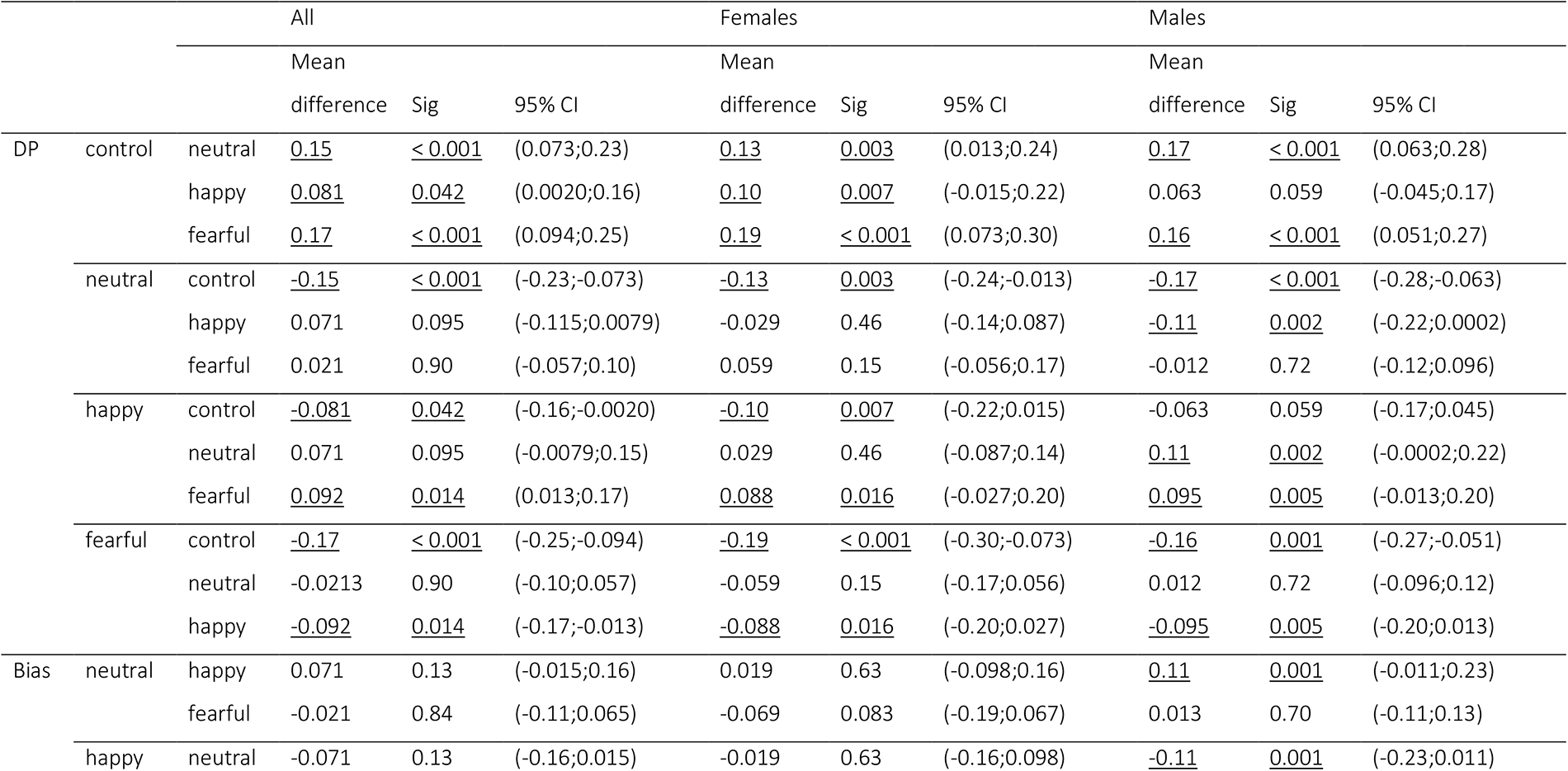
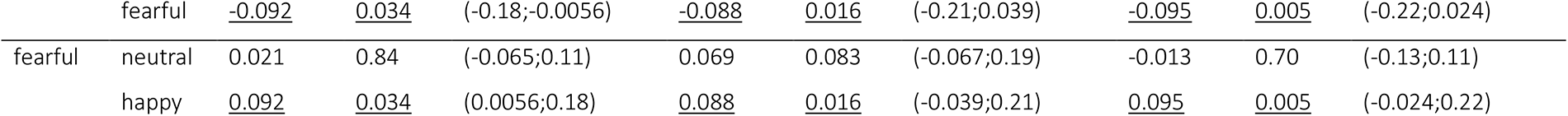
Statistical differences between disengagement probability (DP) and attentional biases toward neutral, happy, or fearful face versus control picture. CI = confidence interval

Females’ attentional bias toward fearful face (vs. control picture, i.e., FE-bias; 0.19, SD 0.27) was significantly higher compared to bias to happy face (vs. control picture, i.e., HA-bias; 0.10, SD 0.26, mean difference 0.088, p = 0.016), while no statistically significant difference was observed between FE-bias and attentional bias to neutral faces (vs. control pictures; NE-bias; 0.13, SD 0.30, mean difference 0.058, p = 0.083). Similarly, males’ FE-bias was significantly higher compared to HA-bias (0.18, SD 0.30 vs 0.06, SD 0.26, mean difference 0.12, p = 0.005). Also, NE-bias (0.17, SD 0.29) was significantly higher compared to HA-bias (mean difference 0.11, p = 0.001) in males. No statistically significant difference was observed between males’ NE-bias and FE-bias (mean difference 0.013, p = 0.70).

### 3.3 Association between attention bias variables and fractional anisotropy

There were no associations between any of the attention biases and WM integrity on the level of the whole sample. We then carried out statistical models in males and females separately. In females, we found negative associations between FE-bias scores and FA values, which were observed in multiple WM regions (including the splenium of corpus callosum (SCC), left anterior limb of internal capsule (ALIC), left posterior limb of internal capsule (PLIC), left posterior thalamic radiation and optic tract (PTR/OR) and left uncinate fasciculus (5000 permutations, p < 0.05)). The result remained significant after controlling for child’s age at scan, maternal pre-pregnancy BMI, maternal age, SES, exposure to smoking, SSRI or SGC during pregnancy and maternal depression (EPDS) or anxiety (SCL-90) scores gathered at 2^nd^ trimester (Figure 1). With postpartum EPDS or SCL-90 scores at child age of 3 months included in the regression model, the result was not statistically significant (TFCE corrected p = 0.08). The associations did not remain statistically significant between FE-bias and mean FA of left uncinate (Spearman ρ = -.052, p = 0.71) and left PTR/OR (Spearman ρ = -.19, p = 0.17) in ROI analyses (Figure 1). No statistically significant associations between males’ FE-bias and FA were observed. Other attentional bias variables showed no associations with FA in the inspected groups.

### 3.4 Association between eye-tracking variables and maternal psychological distress

Self-reported postnatal maternal psychological distress was strongly positively related to females’ FE-bias at five years (Table 4 for correlations in females). EPDS score (r = 0.34, p = 0.010) and SCL-90 score (r = 0.57, p < 0.001) at 3 months postpartum both showed positive association to FE-bias. No significant associations between 2^nd^ trimester EPDS (r = 0.083, p = 0.55) or SCL-90 (r = 0.086, p = 0.55) were observed. In males, no significant correlations between the maternal EPDS or SCL-90 scores were detected (for FE-bias r = -0.020 and p = 0.88 with EDPS at 3 months, and r = 0.11 and p = 0.39 with SCL-90 at 3 months; Supplementary Table 2 and 3 for correlations in males and in all subjects).

**Table 4.** Correlation matrix for females’ eye-tracking results, maternal psychological distress scores and demographical variables with Spearman correlation ρ, p values < 0.05 marked with * and < 0.005 with **. Dp = disengagement probability, CS = control stimulus, FE = fearful, NE = neutral, HA = happy, EPDS = Edinburg postnatal depressive symptoms, SCL-90 = Symptom Checklist, mo = month, trim = trimester, PI = ponderal index, BMI = body mass index, SES = socioeconomic status, SSRI = selective serotonin re-uptake inhibitor, SGC = synthetic glucocorticoid.

|  | dp.<br>1CS | dp.<br>4FE | NE<br>bias | HA<br>bias | FE bias | EPDS<br>3mo | SCL-90<br>3mo | EPDS<br>2nd<br>trim | SCL-90<br>2nd<br>trim | child'<br>s age<br>(year<br>s) | PI | Mate<br>rnal<br>BMI | Mater<br>nal age<br>(years) | smok<br>ing | SSRI | SGC | SES |
| --- | --- | --- | --- | --- | --- | --- | --- | --- | --- | --- | --- | --- | --- | --- | --- | --- | --- |
| dp.1CS | 1.00 |  |  |  |  |  |  |  |  |  |  |  |  |  |  |  |  |
| dp.4FE | 0.09 | 1.00 |  |  |  |  |  |  |  |  |  |  |  |  |  |  |  |
| NE bias | .588** | -0.10 | 1.00 |  |  |  |  |  |  |  |  |  |  |  |  |  |  |
| HA bias | .440** | -0.14 | .472** | 1.00 |  |  |  |  |  |  |  |  |  |  |  |  |  |
| FE bias | .606** | -.710** | .501** | .470** | 1.00 |  |  |  |  |  |  |  |  |  |  |  |  |
| EPDS 3mo | .312* | -0.14 | 0.25 | 0.17 | .343* | 1.00 |  |  |  |  |  |  |  |  |  |  |  |
| SCL-90 3mo | .426** | -.359** | .322* | 0.25 | .574** | .509** | 1.00 |  |  |  |  |  |  |  |  |  |  |
| EPDS 2nd<br>trim | 0.18 | 0.05 | 0.23 | 0.18 | 0.08 | .542** | 0.21 | 1.00 |  |  |  |  |  |  |  |  |  |
| SCL-90 2nd<br>trim | 0.13 | 0.00 | 0.20 | 0.07 | 0.08 | .576** | .512** | .701** | 1.00 |  |  |  |  |  |  |  |  |
| child's age<br>(years) | 0.00 | -0.12 | -0.10 | 0.01 | 0.10 | 0.17 | 0.01 | 0.21 | 0.04 | 1.00 |  |  |  |  |  |  |  |
| PI | -0.01 | -0.04 | 0.05 | -0.08 | 0.02 | 0.10 | 0.12 | -0.07 | 0.10 | -0.04 | 1.00 |  |  |  |  |  |  |
| Maternal<br>BMI | 0.12 | -0.10 | 0.06 | 0.03 | 0.20 | 0.24 | 0.19 | 0.08 | 0.16 | 0.18 | 0.20 | 1.00 |  |  |  |  |  |
| Maternal age<br>(years) | -0.03 | -0.03 | 0.06 | -0.20 | -0.04 | -0.06 | -0.21 | -0.11 | -0.11 | -0.23 | 0.03 | 0.02 | 1.00 |  |  |  |  |
| smoking | 0.20 | 0.18 | 0.23 | 0.11 | -0.02 | 0.14 | 0.13 | -0.15 | -0.05 | -0.17 | 0.12 | 0.20 | 0.18 | 1.00 |  |  |  |
| SSRI | 0.14 | 0.05 | 0.10 | 0.20 | 0.08 | -0.21 | 0.07 | -0.03 | -0.11 | 0.00 | 0.02 | -0.15 | -0.13 | -0.03 | 1.00 |  |  |
| SGC | 0.12 | -0.12 | 0.03 | -0.08 | 0.16 | 0.11 | 0.21 | -0.18 | 0.04 | 0.17 | 0.16 | 0.12 | 0.00 | -0.05 | -0.04 | 1.00 |  |
| SES | 0.00 | 0.18 | -0.02 | 0.06 | -0.13 | 0.09 | 0.00 | 0.17 | 0.07 | 0.04 | -0.09 | -0.06 | 0.24 | -0.18 | 0.15 | -0.12 | 1.00 |

**Figure 1.**
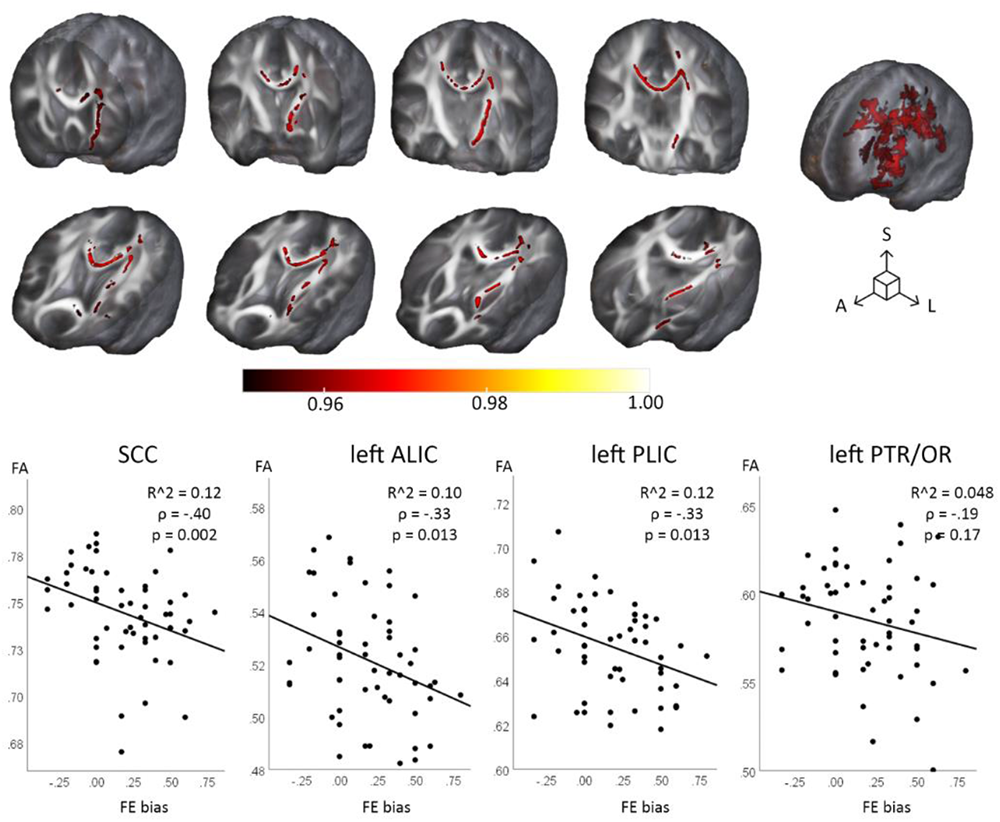
Regions with negative association between FE-bias (fear bias) and fractional anisotropy (FA) value in shown in red. Regression analysis performed with 5000 permutations and threshold-free cluster enhancement correction, p < 0.05. Correlation between FE-bias and FA of entire white matter tract fractional anisotropy (splenium of corpus callosum (SCC), left anterior limb of internal capsule (ALIC), left posterior limb of internal capsule (PLIC), left posterior thalamic radiation and optic tract (PTR/OR)) extracted with co-registration to JHU-ICBM-DTI atlas. A = anterior, S = superior, L = left<colcnt=1>

## 4 Discussion

In the present study, the associations between WM microstructure and attention biases toward emotional facial expressions (happy, fearful, neutral face vs. control, non-face picture) were investigated in a population of typically developing 5-year-old children. We observed that reduced WM integrity (indexed as decrease in FA) in regions of SCC, left uncinate, left ALIC, left PLIC and fronto-occipital tracts predicted higher attentional bias toward fearful expressions in females. Further, maternal postnatal anxiety and depressive symptoms (at 3 months) associated positively with attentional bias toward fearful expression, but only in females. Based on these findings, it is possible that alterations in WM microstructure may transmit long-term effects of maternal mental distress during early life and increase vigilance toward negative emotional expressions.

Faces with emotionally salient expressions have greater tendency to capture attention in a reflexive, involuntary way as compared to neutral faces (Eastwood et al., 2003; E. Fox et al., 2001). This phenomenon is especially well-characterized with negative expressions, signalling potential threat and requiring immediate actions (Eastwood et al., 2003; E. Fox et al., 2001; Surguladze et al., 2003; Vuilleumier & Schwartz, 2001). In addition, we observed 5-year-olds to disengage from control stimulus faster compared to faces with neutral, happy, or fearful expressions, and repeated similar pattern previously detected with infants (E. L. Kataja et al., 2020). The emotional valence of the face has been detected to enhance the activation of FFA, suggesting the existence of feedback connections between the amygdala and visual cortex (Ishai et al., 2004; Vuilleumier & Pourtois, 2007). This top-down control can guide stronger engagement of attention when more salient facial stimuli are present. Higher negative reactivity at the age of three years was also indicated to associate with higher activation in both the amygdala and FFA, but in contrast, lower connectivity between the areas during face processing. The activation patterns of face-selective regions reach adult-like functional specificity only during late adolescence (Cohen Kadosh et al., 2011; Scherf et al., 2007), even though the activity of FFA and the core face network is detected already at age of 7 years (Cantlon et al., 2011; Cohen Kadosh et al., 2011). The extended face network in children presents hyperactivation compared to selective activation in adults suggesting that the task-specific engagement of regions is maturated during development (Haist et al., 2013).

The accuracy of recognizing different facial emotional expressions is improving during childhood (Chronaki et al., 2015; C. Herba & Phillips, 2004). Happiness is the emotion recognized earliest and with the most accuracy during childhood when assessed in comparison to detection of facial expressions of sadness, fear, angriness, and disgust (Gao & Maurer, 2010; Mancini et al., 2013; Rodger et al., 2015). In our study, the children disengaged fastest from the happy face toward distractor, which might indicate them to recognise the expression fastest and to be ready to move forward. On the other hand, sadness is the emotional state with highest misidentification rates and slowest improvement of recognition during childhood from the age of four years to adolescence (Chronaki et al., 2015; Gao & Maurer, 2010; C. Herba & Phillips, 2004). The identification of neutral facial expression has been suggested to develop later between ages from 8 to 11 years (Mancini et al., 2013). Apart from improvement of recognizing the emotional state of other individuals, the affective reactions to experienced emotions maturate during childhood (Mancini et al., 2013). The emotional arousal related to emotional facial expressions decreases while the child develops more mature ways of processing emotional information.

We observed maternal early postpartum depressive and anxiety symptoms to predict attentional bias to fearful faces at five years of age, but only in females. Multiple previous studies have shown females to be especially sensitive to maternal psychological distress (Braithwaite et al., 2016; Dean et al., 2018; Erickson et al., 2019; Quarini et al., 2016; Simcock et al., 2016; Wen et al., 2017), and that brain development shows sexual dimorphism in both normal growth and in response to environmental exposures (Hashempour et al., 2023; Kumpulainen, Merisaari, et al., 2023; Lautarescu et al., 2020; Lehtola et al., 2020, 2022). In a prior study of emotional attention, researchers found that daughters of depressed mothers attended selectively to sad faces, while controls and sons of depressed mothers did not (Kujawa et al., 2011). In addition, our own study showed that maternal depression during the pre-or postpartum periods associated with higher fear bias in 8-month-old infants (E.-L. Kataja et al., 2020), and maternal anxiety symptoms during the same periods enhanced female infants’ attention towards all faces whereas it decreased males’ attention (E.-L. Kataja et al., 2019). Our current findings further consolidate these conclusions of sex-specific sensitivity to early exposure to internalizing symptoms which is detectable across childhood attention patterns. Maternal negative behaviour during a problem-solving interaction has also been associated with increased amygdala activation during processing of angry and fearful faces (Pozzi et al., 2020). Previous studies (see Table 1 for literature review) have shown children’s attentional bias to threat signals to predict irritability in 6-14-year-olds (Salum et al., 2017) and overall anxiety in 13-year-olds (Abend et al., 2018). Anxious children were also faster to detect fear (Simcock et al., 2020) and showed stronger overall attention to affective stimuli (Waters et al., 2004) between ages of 9 and 12 years. Behavioural inhibition as toddlers was also associated with later increased attention to threat (Pérez-Edgar et al., 2010), and further, in another study of 5-7-year-old children (White et al., 2017), behavioural inhibition was observed to predict anxiety symptoms in children with attentional bias to threat. In some studies, attentional avoidance of negative emotions was associated to anxiety symptoms between ages 9 and 13 years (Brown et al., 2013; Kallen et al., 2007), and in relation to early life stress (Humphreys et al., 2016). In addition, Waters et al. (2004) observed stronger attentional bias to fear-related pictures in females compared to males, while no sex-differences were detected in the other studies.

The amygdala and limbic system connectivity to widely distributed structures (insula, cingulate gyrus ang prefrontal cortex) goes over remarkable alterations in emotion-related brain circuitry during early childhood, strongly suggesting it has a role in the emotional maturation during first years of life and is especially susceptible to early life adversity (Gee et al., 2013; Qin et al., 2012). The reciprocal connections between the amygdala and different regions of PFC, for example OFC and ventromedial prefrontal cortex (vmPFC), have been shown to participate in reappraisal of negative emotions (Buhle et al., 2014; Wager et al., 2008). The downregulation of the amygdala activation during reappraisal after emotionally relevant stimuli has been shown to be enhanced with age (Pitskel et al., 2011). Uncinate fasciculus connects amygdala and prefrontal cortex and is kept as a crucial link in emotion processing and regulation (Swartz et al., 2015), but WM fibers connecting prefrontal cortex, amygdala, hippocampus, and thalamus run also along anterior internal capsule (Coenen et al., 2020; Safadi et al., 2018). Besides the amygdala, the bed nucleus of the stria terminalis (BNST), has been indicated to play a part in directing attention to threatening cues (Herrmann et al., 2016; Somerville et al., 2010; Walker et al., 2003). Furthermore, the BNST is involved in the emotional face perception (Sladky et al., 2018). Both the amygdala and the BNST have been shown to activate in response to threatening signals, but with slightly separate functional roles. The amygdala shows engagement in situations with explicit and expected threat, while the BNST seems to respond preferentially to more ambiguous threat signals (Dzafic et al., 2019; Naaz et al., 2019) and anticipation of threat, and its sustained activation is associated with hyperarousal and hypervigilance (Chang et al., 2016). The integrity of stria terminalis, connecting these regions of extended amygdala network, correlates positively with faster recognition of anger with or without prior cues (Dzafic et al., 2019). Besides the amygdala, BNST shows also tight connections to PFC, caudate and thalamus (Klumpers et al., 2017).

Considering the complexity of the brain circuitry involved, it is unsurprising that widespread alterations of WM microstructure are associated with impairments and problems with emotion processing. We observed that increased attention toward fearful facial expression was associated with reduced WM integrity in multiple tracts, mainly in left hemisphere. The immaturity of especially fronto-thalamic and fronto-limbic tracts like uncinate and anterior internal capsule might imply weaker connections and downregulation of limbic area activation in response to emotional stimuli. The connectivity between OFC and the amygdala has been indicated as essential for decreasing negative affect (Banks et al., 2007) and higher activity of the amygdala is associated with negative emotionality (e.g. tendency to experience and react with negative emotions (Everaerd et al., 2015; Haas et al., 2007; Stein et al., 2007)). In a previous study, age-related decline in the amygdala activation to emotional faces was associated with increased FA within the uncinate fasciculus in children and adolescents in the left hemisphere, and on the other hand, the increased amygdala activation to sad faces was related to greater internalizing symptoms (Swartz et al., 2015). Furthermore, increased amygdala reactivity during processing of emotional stimuli is associated with anxiety related temperamental features (Stein et al., 2007). Reduced FA of uncinate has been previously associated with anxiety (Adluru et al., 2017; Kaczkurkin et al., 2016; Liao et al., 2014; Lichtin et al., 2021; Tromp et al., 2019) and depression (Cullen et al., 2010; Huang et al., 2006; LeWinn et al., 2014; Mohamed Ali et al., 2019) together with reduced FA of ALIC (Bessette et al., 2014; Ghazi Sherbaf et al., 2018; Henderson et al., 2013) in children and adolescents. Based on these prior findings, we propose that our results imply that increased attentional bias to fear might culminate as an increased risk for negative emotionality and internalizing symptoms with inferior capacity to reappraise and suppress negative emotions, and this interaction is transmitted by decreased WM integrity in left-sided fronto-limbic and fronto-thalamic tracts.

The current study has some noteworthy limitations. The sample size is acknowledged to be moderate, especially when females and males are examined separately. Henceforth, the research needs to be replicated with larger populations to ensure their repeatability. Related to eye-tracking measurements, use of mobile/interactive figures or familiar faces might have increased the sensitivity for recognition of emotional expressions; however, this is methodologically challenging as the stimulus need to be uniform across population. Furthermore, maternal psychological distress scores were overall quite low in this study population, which may have lead part of their associations to remain unnoticed.

## 5 Conclusion

We were able to make novel discoveries on the links between fear bias and the brain WM integrity that were specific for females. Further, this association was dependent on maternal early postnatal depressive symptoms, again only in females. Future studies are needed for increasing our understanding on the practical relevance of these findings and the role that the observed sexual dimorphism has later in development. Based on our findings, it is possible that alterations in WM microstructure may transmit the long-term effects of maternal mental distress during early life and increased vigilance toward negative emotional expressions or that these phenotypes share common developmental pathways that are sensitive to maternal perinatal distress.

## Acknowledgements

We thank our research nurse Susanne Sinisalo for her expertise in study management and performing the scans with the investigators and all participating FinnBrain families. We would also like to thank John D. Lewis for help planning the DTI sequences, Harri Merisaari for developing the DTI processing pipelines, Riitta Parkkola for screening the MRI for incidental findings, Tuire Lähdesmäki for being the consulting child neurologist for the studies, and Jukka M. Leppänen for helping with the eye-tracking data pre-processing.

## Author contributions

- VK: participated in recruiting children to MRI sub study, gathering MRI data, pre-processing and analysing DTI data, performing statistical analyses and writing the manuscript
- ELK: participated in gathering, pre-processing and analysing eye-tracking data and writing the manuscript
- AC, ESi and EPP: participated in recruiting children to MRI sub study, gathering MRI data and writing the manuscript
- ESa: gathering MRI data
- LK and HK: funding acquisition, designing and establishing the FinnBrain Birth cohort, building infrastructure for carrying out the study
- JJT: study conception, funding acquisition, supervision of VK, planned and implemented the MRI acquisition parameters, building the DTI pre-processing pipelines, pre-processing and analysing eye-tracking data, planned the analytical approach and performed the data analyses.
- TH: eye-tracking methodology

## Data availability statement

The Finnish law and our ethical permissions do not allow openly sharing of the data used in this study. Investigators who are interested in getting access to the data can contact FinnBrain Birth Cohort administration (https://sites.utu.fi/finnbrain/en/contact/).

