## Supplementary Figure 1, Tables 1-3 for "Attentional bias toward fearful faces is associated with maternal postnatal distress and alterations in white matter microstructure in 5-year-old females"

Supplement Figure 1.

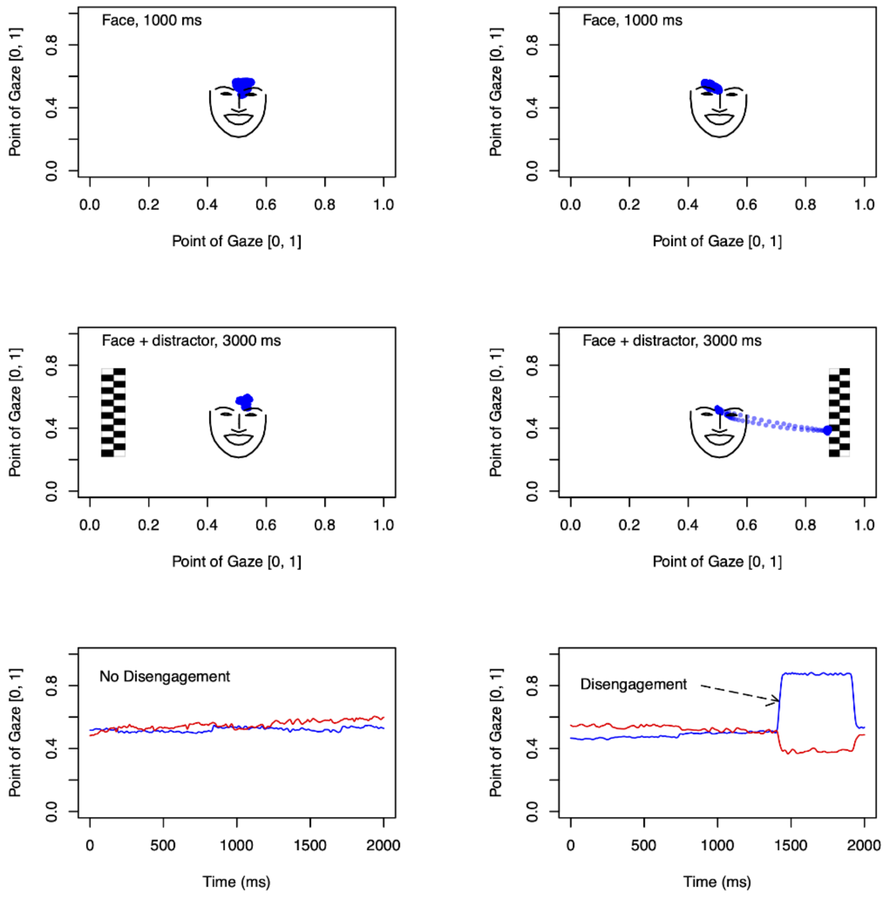


*Note*. Participants were presented with a face or a non-face in the center of the screen. One second later, a "distractor" was added to the left or to the right.  Left: An example of a "no disengagement" trial in which the gaze does not shift from the central to the lateral stimulus. Right: An example of a "disengagement" trial in which the gaze shifts rapidly from the central to the lateral stimulus. The X- and Y-coordinates of the point of gaze on the display are shown by the blue and red lines, respectively. The outline and key landmarks of the face stimulus are shown based on values extracted by the OpenFace 2.2.0 toolkit (Baltrusaitis et al., 2018).

Supplement Table 1. Eye-tracking variable names and explanations.

| Eye-tracking variables | |
|---|---|
| Dp.1CS | Probability of disengagement from control picture toward a distractor stimulus |
| Dp.2NE | Probability of disengagement from neutral face picture toward a distractor stimulus |
| Dp.3HA | Probability of disengagement from happy face picture toward a distractor stimulus |
| Dp.4FE | Probability of disengagement from fearful face picture toward a distractor stimulus |
| NE-bias | Attentional bias toward neutral face versus control picture |
| HA-bias | Attentional bias toward happy face versus control picture |
| FE-bias | Attentional bias toward fearful face versus control picture |
| Mean-DP-face | Average probability of disengagement from faces (neutral/happy/fearful) |

Supplement Table 2. Correlation matrix for eye-tracking results, maternal psychological distress scores and demographical variables in boys with Spearman correlation ρ, p values < 0.05 marked with * and < 0.005 with **. Dp = disengagement probability, CS = control stimulus, FE = fearful, NE = neutral, HA = happy, EPDS = Edinburg postnatal depressive symptoms, SCL-90 = Symptom Checklist, mo = month, trim = trimester, PI = ponderal index, BMI = body mass index, SES = socioeconomic status, SSRI = selective serotonin re-uptake inhibitor, SGC = synthetic glucocorticoid.

| | dp. 1CS | dp. 4FE | NE bias | HA bias | FE bias | EPDS 3mo | SCL-90 3mo | EPDS 2nd trim | SCL-90 2nd trim | child's age (years) | PI | Maternal BMI | Maternal age (years) | smoking | SSRI | SGC | SES |
|---|---|---|---|---|---|---|---|---|---|---|---|---|---|---|---|---|---|
| dp.1CS | 1.000 | | | | | | | | | | | | | | | | |
| dp.4FE | 0.165 | 1.000 | | | | | | | | | | | | | | | |
| NE bias | .357$^{**}$ | -.313$^{*}$ | 1.000 | | | | | | | | | | | | | | |
| HA bias | .449$^{**}$ | -.280$^{*}$ | .517$^{**}$ | 1.000 | | | | | | | | | | | | | |
| FE bias | .450$^{**}$ | -.753$^{**}$ | .604$^{**}$ | .545$^{**}$ | 1.000 | | | | | | | | | | | | |
| EPDS 3mo | 0.001 | 0.004 | -0.029 | 0.127 | -0.020 | 1.000 | | | | | | | | | | | |
| SCL 3mo | -0.040 | -0.089 | 0.101 | 0.107 | 0.061 | .563$^{**}$ | 1.000 | | | | | | | | | | |
| EPDS 2nd trim | 0.117 | 0.023 | 0.003 | 0.204 | 0.060 | .566$^{**}$ | .547$^{**}$ | 1.000 | | | | | | | | | |
| SCL 2nd trim | 0.080 | -0.077 | 0.081 | .262$^{*}$ | 0.114 | .638$^{**}$ | .621$^{**}$ | .637$^{**}$ | 1.000 | | | | | | | | |
| child's age (years) | -0.099 | 0.016 | -0.208 | -0.201 | -0.124 | -0.058 | -0.089 | -0.085 | -0.171 | 1.000 | | | | | | | |
| PI | -0.061 | -0.118 | 0.041 | -0.064 | 0.009 | -0.025 | -0.076 | -0.249 | -0.122 | 0.177 | 1.000 | | | | | | |
| Maternal BMI | -0.175 | 0.038 | -0.080 | -0.102 | -0.156 | 0.076 | 0.163 | -0.123 | 0.067 | -0.014 | .414$^{**}$ | 1.000 | | | | | |
| Maternal age (years) | -0.085 | -0.109 | 0.031 | 0.115 | 0.086 | 0.072 | -0.089 | 0.128 | 0.078 | 0.058 | -0.018 | -0.120 | 1.000 | | | | |
| smoking | -0.047 | 0.086 | 0.159 | -0.086 | -0.041 | -0.022 | 0.161 | -0.240 | -0.096 | 0.131 | 0.018 | .278$^{*}$ | -.405$^{**}$ | 1.000 | | | |
| SSRI | 0.018 | -0.045 | 0.050 | 0.129 | 0.080 | 0.261 | 0.124 | 0.064 | 0.184 | 0.223 | -0.244 | -0.069 | -0.077 | -0.061 | 1.000 | | |
| SGC | -0.178 | .276$^{*}$ | -0.135 | -0.192 | -.306$^{*}$ | 0.008 | 0.045 | -0.050 | -0.005 | 0.133 | 0.005 | 0.049 | 0.120 | 0.111 | 0.199 | 1.000 | |
| SES | 0.031 | -0.131 | 0.109 | 0.095 | 0.150 | -0.173 | -0.068 | 0.015 | 0.095 | -0.011 | 0.180 | 0.025 | 0.217 | -.258$^{*}$ | -0.257 | -.352$^{**}$ | 1.000 |

Supplement Table 3. Correlation matrix for eye-tracking results, maternal psychological distress scores and demographical variables in all subjects with Spearman correlation ρ, p values < 0.05 marked with * and < 0.005 with **. Dp = disengagement probability, CS = control stimulus, FE = fearful, NE = neutral, HA = happy, EPDS = Edinburg postnatal depressive symptoms, SCL-90 = Symptom Checklist, mo = month, trim = trimester, PI = ponderal index, BMI = body mass index, SES = socioeconomic status, SSRI = selective serotonin re-uptake inhibitor, SGC = synthetic glucocorticoid.

| | dp. 1CS | dp. 4FE | NE bias | HA bias | FE bias | EPDS 3mo | SCL-90 3mo | EPDS 2nd trim | SCL-90 2nd trim | child's age (years) | PI | Mater nal BMI | Mater nal age (years) | smoki ng | SSRI | SGC | SES |
|---|---|---|---|---|---|---|---|---|---|---|---|---|---|---|---|---|---|
| dp.1CS | 1.000 | | | | | | | | | | | | | | | | |
| dp.4FE | 0.151 | 1.000 | | | | | | | | | | | | | | | |
| NE bias | .478** | -.200* | 1.000 | | | | | | | | | | | | | | |
| HA bias | .440** | -.216* | .487** | 1.000 | | | | | | | | | | | | | |
| FE bias | .525** | -.723** | .541** | .506** | 1.000 | | | | | | | | | | | | |
| EPDS 3mo | 0.157 | -0.050 | 0.085 | 0.150 | 0.154 | 1.000 | | | | | | | | | | | |
| SCL 3mo | .194* | -.214* | .192* | 0.174 | .328** | .524** | 1.000 | | | | | | | | | | |
| EPDS 2nd trim | 0.150 | 0.017 | 0.104 | .208* | 0.082 | .557** | .386** | 1.000 | | | | | | | | | |
| SCL 2nd trim | 0.105 | -0.049 | 0.128 | 0.170 | 0.110 | .612** | .573** | .669** | 1.000 | | | | | | | | |
| child's age (years) | -0.052 | -0.046 | -0.177 | -0.097 | -0.026 | 0.037 | -0.035 | 0.060 | -0.060 | 1.000 | | | | | | | |
| PI | -0.036 | -0.077 | 0.035 | -0.082 | 0.017 | 0.020 | 0.025 | -0.170 | -0.021 | 0.070 | 1.000 | | | | | | |
| Maternal BMI | -0.041 | -0.009 | -0.027 | -0.044 | -0.002 | 0.150 | 0.181 | -0.003 | 0.114 | 0.070 | .311** | 1.000 | | | | | |
| Maternal age (years) | -0.048 | -0.058 | 0.049 | -0.042 | 0.018 | -0.011 | -0.129 | -0.017 | -0.010 | -0.099 | 0.006 | -0.083 | 1.000 | | | | |
| smoking | 0.064 | 0.127 | .192* | -0.005 | -0.036 | 0.042 | 0.143 | -.190* | -0.077 | 0.008 | 0.051 | .251** | -.186* | 1.000 | | | |
| SSRI | 0.067 | -0.008 | 0.079 | 0.147 | 0.075 | 0.045 | 0.105 | 0.021 | 0.068 | 0.133 | -0.141 | -0.108 | -0.087 | -0.044 | 1.000 | | |
| SGC | -0.043 | 0.123 | -0.058 | -0.141 | -0.105 | 0.054 | 0.123 | -0.123 | 0.014 | 0.143 | 0.075 | 0.086 | 0.067 | 0.057 | 0.118 | 1.000 | |
| SES | 0.017 | 0.007 | 0.046 | 0.074 | 0.011 | -0.030 | -0.036 | 0.096 | 0.083 | 0.010 | 0.057 | -0.013 | .250** | -.221* | -0.093 | -.247** | 1.000 |
